# Far-Red Light in Early Growth Stages Boosts Lettuce Biomass and Preserves Anthocyanins

**DOI:** 10.1101/2025.07.22.666241

**Authors:** Christopher P. Levine, Keiichiro Tanigawa, Yu Wakabayashi, Wei Guo, Yuchen Qu, Ichiro Terashima, Wataru Yamori

## Abstract

**Background and Aims:** Light plays a dual role in plants, serving as both an energy source and a regulator of development from seedling to senescence. Recently, far-red (FR) radiation has gained attention in the controlled environment agriculture (CEA) science and grower community for its potential to enhance yield through canopy expansion and improved light capture, contributing positively to photosynthesis. This study explores how supplemental FR light promotes lettuce growth and morphology across weekly intervals as well as analyzing photosynthetic parameters, pigment accumulation, and anthocyanin gene expression.

**Methods:** Red leaf lettuce (*Lactuca sativa* ‘Red Fire’) was grown in a commercial plant factory with artificial light for six weeks. 5000K white (W) light was maintained at 300 μmol m^-2^ s^-1^, and FR, when supplemented, was added at 100 μmol m^-2^ s^-1^ in addition to the 300 μmol m^-2^ s^-1^ of W light. Four lighting treatments were tested under a 16 h photoperiod: (1) W for all 6 weeks (“W”), (2) 4 weeks of W followed by 2 weeks of supplemental FR (“W to W+FR”), (3) 4 weeks of FR supplementation followed by 2 weeks of only W (“W+FR to W”), and (4) W+FR for all 6 weeks (“W+FR”).

**Key Results:** The shoot dry weight after 6 weeks in “W+FR”, “W+FR to W” and “W to W+FR” was greater than “W”. Both “W+FR” and “W+FR to W” showed a tendency for greater canopy expansion compared to “W” as well as “W to W+FR”. There were no significant differences in stomatal conductance among the treatments. On the other hand, in both “W” and “W+FR to W” plants, CO_2_ assimilation rates were enhanced when FR light was supplemented during measurement, compared to when FR was not provided. Anthocyanin accumulation was greater in both “W” and “W+FR to W”, consistent with the expression of key genes involved in the anthocyanin biosynthesis pathway, including anthocyanin synthase (*ANS*), flavanone 3-hydroxylase (*F3H*) and dihydroflavonol 4-reductase (*DFR*).

**Conclusions:** This study demonstrates that FR supplementation during the early growth stages of lettuce promotes biomass accumulation by enhancing both canopy expansion and photosynthetic activity, while maintaining high levels of functional compounds such as anthocyanins.

## Introduction

The global food system accounts for 30% of annual greenhouse gas emissions, with 19% attributed to food transport, particularly for refrigerated fruits and vegetables (Li et al., 2022). Local crop production currently fulfills less than one-third of global food demand (Li et al., 2022; Kinnunen et al., 2020). Mitigating environmental impact requires prioritizing local food production, particularly in affluent nations, where food-mile emissions are a significant contributor. Controlled environment agriculture (CEA) presents a promising approach for enhancing local food self-sufficiency, especially in light of supply chain disruptions caused by geopolitical instability and global crises such as pandemics (Li et al. 2025; Furuta et al. 2025). However, CEA faces several obstacles, including challenges related to profitability, energy efficiency, supportive public policy, and consumer acceptance (van Delden et al., 2021).

Supplementing plant lighting with far-red (FR) radiation has gained considerable attention in recent years. It is well known that FR induces shade avoidance responses, leading to stem elongation, while shade tolerance mechanisms promote leaf expansion in various but not all plants (Zhen et al., 2020; Kusuma et al., 2023; Owen et al., 2015; Van de Velde et al., 2023; Skabelund et al., 2025). Studies indicate that FR photon supplementation enhances plant biomass accumulation by promoting leaf expansion, leading to increased light capture (Meng and Runkle, 2019; Li and Kubota, 2009; Park and Runkle, 2017; Zhen et al., 2020).

Exposure to FR light triggers the conversion of phytochromes from their active (Pfr) to inactive (Pr) forms, with peak absorption occurring at 730 nm. In contrast, red light (670 nm) reverts phytochromes to their active state (Sharrock, 2008; Kaiser et al., 2024). Notably, the photon flux density (PFD) of FR photons (∼700-750 nm) is 18-19% of the photosynthetic photon flux density (400-700 nm) under direct unfiltered sunlight (ASTM standard, 2020). However, most horticultural LED fixtures for CEA emit less than 2% as FR except for some lights designed specifically to regulate flowering (Runkle, 2022).

Furthermore, FR supplementation has also been found to reduce anthocyanin concentration in lettuce cultivars, which may affect consumer purchasing preferences (Van Brenk et al., 2024). Additionally, higher FR intensities in choy sum (*Brassica rapa* subsp*. chinensis* var*. parachinensis*) have been linked to a decrease in total carotenoid content (Zou et al., 2024), while in red butterhead lettuce, FR supplementation significantly reduced concentrations of calcium, magnesium, manganese, and phosphorus (Van Brenk et al., 2024). While FR radiation can enhance biomass accumulation, it may cause dilution effects— reducing concentrations of certain compounds per unit leaf area or mass without lowering total contents per plant. For instance, Kelly and Runkle (2024) found that partial substitution of red with FR light increased leaf area, diluting anthocyanin concentrations and pigmentation. This highlights the need to consider measurement units and balance trade-offs when applying FR effectively.

Beyond mineral and phytochemical accumulation, another critical aspect influencing the effects of FR on plant growth is the practical context of dense growing environments, where overlapping leaves play a major role. Zhen and Bugbee (2020) reported equivalent canopy photosynthesis with partial FR substitution across 14 species. However, their plants were grown and acclimated in a glass-covered greenhouse under unspecified high or low light treatments, from which subsequent short-term photosynthetic parameters were collected. Since plant photosynthetic responses can vary with light acclimation, particularly under prolonged exposure, the generalization of these results to plant factories with 100% artificial light is uncertain. Notably, Jeong et al. (2024) found that 20% FR substitution significantly reduced net photosynthesis in lettuce leaves acclimated to FR, and Zou et al. (2019) observed significant declines in photosynthetic rates after 14–17 days of FR acclimation.

Overall, the objective of this research is to evaluate FR radiation supplementation under practical commercial plant factory with artificial light (PFAL) conditions. Specifically, we assess the effects of FR supplementation on a red leaf lettuce cultivar exposed to a background photosynthetic photon flux density (PPFD) of 300 μmol m^-2^ s^-1^ 5000 K white light over six weeks. Although Zhen and Bugbee (2020) claim that PPFD (400–700 nm) should be substituted with FR to achieve equal canopy quantum yield for CO_2_ fixation, we supplemented FR without changing the PPFD (400–700 nm); the rationale for this approach will be discussed in the discussion section. Parameters of interest focused on yield, morphology, plant architecture, anthocyanin gene expression, stomatal conductance, photosynthesis, and metabolite profiles across two defined treatment phases, with weekly measurements capturing morphological changes over time. We hypothesize that supplemental FR radiation during the early growth stages (starting at beginning of week 1 through end of week 4) increases lettuce biomass via canopy expansion and photosynthetic performance, without reducing anthocyanin accumulation.

## Material and Methods

### Plant materials and experimental setup

The experiment was conducted in a commercial PFAL in Tanashi, Nishitokyo, Japan (Plants Laboratory Inc., Tokyo, Japan). The pelleted seeds of ‘Red Fire’ red leaf lettuce (*Lactuca sativa*, Takii Seed Co., Kyoto, Japan) were used. Urethane cubes were hydrated with reverse osmosis water and hydroponic fertilizer to an electrical conductivity of 1.50 ± 0.05 dS m^−1^ (GG liquid A & B stock solutions, Green Green Co., Ltd, Fukuoka, Japan). The environmental conditions in the PFAL were controlled and logged in real time, maintaining an air temperature of 22 ± 2 °C (day/night), a relative humidity (RH) of 63 ± 6%, and a 16 h photoperiod. 5000K white LED lights (TecoG II-40N2-5-23, Toshin Electric Co., Ltd., Osaka, Japan) were used along with FR LED lights (NK system 11A-Z20E4073, Nippon Medical and Chemical Instruments Co., Ltd., Osaka, Japan; see the wavelength in Fig. 1B, 1C). The 4 lighting treatments include 1) 6 weeks of white light (control) (W), 2) 4 weeks of white light followed by 2 weeks of FR light (W to W+FR), 3) FR light for 4 weeks followed by 2 weeks of white light (W+FR to W), and 4) continuous FR light for the entire 6 weeks (W+FR) (Fig 1A). The first 4 weeks only had 2 treatments which consisted of only (W) and (W+FR). Starting from week 5, we had (W), (W+FR), (W to W+FR), and (W+FR to W) treatments which is why we only had treatment 1 (W) and 4 (W+FR) for the first 4 weeks in Fig 3. The 4-week transplant point reflects standard practice used in the operational system of an established commercial head lettuce PFAL in Japan, where transplanting from the second-highest density is required due to plant growth canopy expansion. While the 4-week transplant point aligned well with our system’s specific combination of cultivar, lighting treatments, temperature setpoints, and tray densities, we acknowledge that different cultivars or system configurations may warrant alternative transplant timings. 5000K white light was provided at 300 μmol m^−2^ s^−1^, and when additional FR was supplemented, it was applied at 100 μmol m^−2^ s^−1^ (Fig. 1C, 1D). More detailed light description is provided in Table 1. The light was thoroughly mapped and measured from the base of the growing area to ensure consistency between treatments.

**Figure 1.**
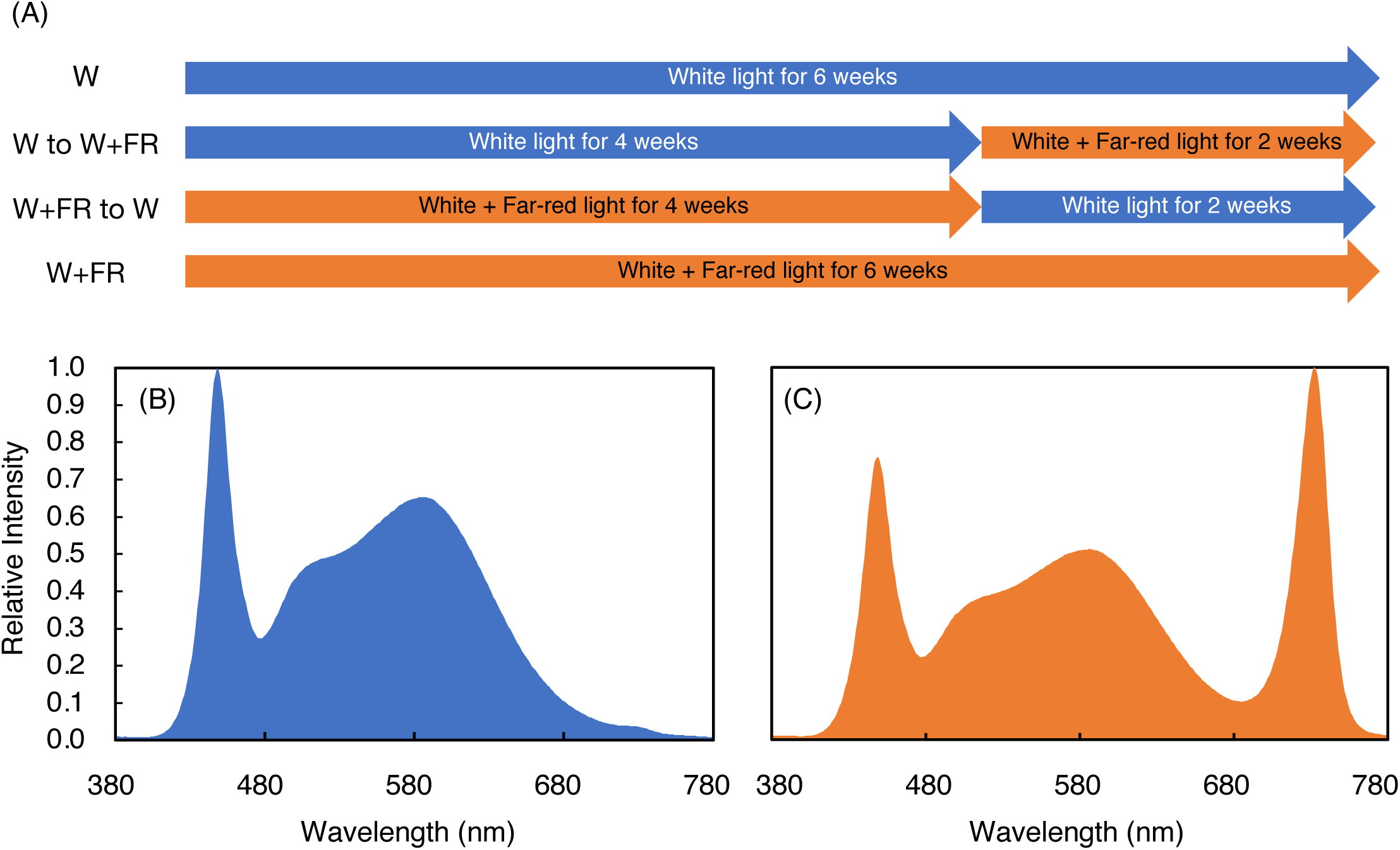
(A) The four experimental light treatments, each consisting of a 16-hour photoperiod followed by an 8-hour dark period: (1) white (W) light only for 6 weeks as the control, (2) 4 weeks of W light followed by 2 weeks of supplemental far-red (FR) (W to W+FR), (3) 4 weeks of FR supplementation followed by 2 weeks of W light (W+FR to W), and (4) FR supplementation for 6 weeks (W+FR). (B) Representative spectral photon distributions of W lighting treatments; 5000k white LED containing 300 µmol m^−2^ s^−1^ of 400-700 nm photosynthetically active radiation (PAR). (D) W+FR treatment containing 300 µmol m^−2^ s^−1^ PAR and 100 µmol m^−2^ s^−1^ of FR photons.

**Table 1.** Spectral characteristics of W and W+FR lighting treatments delivered by 5000k white and FR LEDs.

| Integrated photon flux density ($\mu\text{mol m}^{-2} \text{s}^{-1}$) | | W | W+FR |
| --- | --- | --- | --- |
| PPFD (Photosynthetic photon flux density) | 400–700 nm | 299 | 301 |
| FR-PFD (Far-Red photon flux density) | 700–750 nm | 5 | 84 |
| TPFD (Total photon flux density) | 400–750 nm | 303 | 385 |
| YPFD (Yield photon flux density) <sup>1</sup> | 380–780 nm | 255 | 270 |
| **Daily light integral ($\text{mol m}^{-2} \text{day}^{-1}$)** |  |  |  |
| PAR DLI <sup>2</sup> | 400–700 nm | 17.2 | 17.4 |
| FR DLI <sup>3</sup> | 700–750 nm | 0.3 | 4.8 |
| TPFD DLI <sup>4</sup> | 400–750 nm | 17.5 | 22.2 |
| YPFD DLI <sup>5</sup> | 380–780 nm | 14.7 | 15.6 |
| **Waveband % of TPFD** |  |  |  |
| Blue | 400–500 nm | 29% | 22.9% |
| Green | 500–600 nm | 45.7% | 35.6% |
| Red | 600–700 nm | 23.9% | 19.7% |
| Far-Red | 700–750 nm | 1.5% | 21.8% |
| **Waveband % of PPFD** |  |  |  |
| Blue | 400–500 nm | 29.4% | 29.2% |
| Green | 500–600 nm | 46.3% | 45.4% |
| Red | 600–700 nm | 24.3% | 25.3% |
| Far-Red | 700–750 nm | 1.5% | 27.6% |

<sup>1</sup> YPFD ( $\mu\text{mol m}^{-2} \text{s}^{-1}$ , 380–780 nm): Biologically weighted photon flux calculated as $\text{YPFD} = \sum[\lambda=380 \text{ to } 780] \text{SPD}(\lambda) \times \text{RQE}(\lambda)$ where SPD is the spectral photon flux density ( $\mu\text{mol m}^{-2} \text{s}^{-1} \text{nm}^{-1}$ ) and RQE is the relative quantum efficiency. The RQE spectrum is from Sager et al. (1988), originally tabulated at 2 nm intervals and linearly interpolated to 1 nm resolution to match the measurement data. This metric reflects photosynthetically weighted photon flux from 380 nm to 780 nm. <sup>2</sup> PPFD × 16 h photoperiod. <sup>3</sup> FR PFD × 16 h photoperiod. <sup>4</sup> TPFD × 16 h photoperiod. <sup>5</sup> YPFD × 16 h photoperiod.

These densities were selected primarily to emulate production practices of a commercial PFAL in Japan. During the first 2 weeks of the experiment, 300 plants for each treatment were grown in a urethane cube mat at a density of 1847 plants per m^2^ (Fig. S1). After 2 weeks, 200 healthy plants were randomly selected and transplanted into a custom-built nutrient film technique (NFT) system for weeks 3 and 4 at a density of 314 plants per m^2^ (Fig. S1). The NFT system used the same reverse osmosis water, hydroponic fertilizer, and concentration. After 4 weeks, 60 healthy plants were randomly selected and transplanted into a similar NFT system for weeks 5 and 6 except at a density of 63 plants per m^2^ (Fig. S1).

### Plant Growth Analysis

Each experiment run began with 300 seeds planted per treatment. Every week for a total of 6 weeks, 5 to 10 plants were randomly selected and harvested for destructive measurements. To minimize edge effects, only plants surrounded by neighboring plants were used for data collection. Border and corner plants were excluded due to increased exposure to possible airflow and microclimatic variability, which can alter growth and make them less representative of the overall treatment. Plants were measured for total shoot fresh mass, leaf area, and finally dried for 1 week at 80°C in a constant-temperature oven for dry shoot mass measurement. We determined leaf surface area values using a LI-3100C Area Meter (LI-COR Biosciences, Lincoln, NE, USA), and all shoot biomass was used to measure average leaf surface area and specific leaf area. Every effort was made to flatten leaf tissue as much as possible during leaf surface area measurements.

For the plant shape analysis, plants were grown under the same experimental conditions. The main axis, a plant height parameter (taken from side images), and canopy equivalent diameter were measured (Fig. 5B).

Once a week, five randomly selected and consistently tagged plants were temporarily removed, and side and top-view images were captured with plants placed inside a 64 × 64 × 64 cm white photography box illuminated with white LED light. Images were captured using an iPhone 12 Pro Max (Apple Inc., Cupertino, CA, USA) at 1× (no zoom), with the camera focused on the lettuce plant and positioned at a fixed distance of approximately 70 cm from the center of the urethane propagation cube where the seed was inserted. Plants were imaged without any structural support; therefore, the recorded plant height reflects the natural height under gravitational constraints, not the maximum extended length. From the side images, plant height (main axis length) was measured, while top-view images were used to calculate canopy equivalent diameter. Plant shape characteristics which include top surface area, side surface area, canopy coverage (also referred to as “extent” in MathWorks, 2024), main axis length, and canopy equivalent diameter were analyzed using EasyPCC_V2 software (Guo et al., 2017), which applies a decision-tree-based segmentation model to accurately extract vegetation features from images under varied lighting conditions.

The canopy equivalent diameter of the circle (Fig. 5A) is of the same area as the pixel surface area of vegetative tissue from the top canopy using the following equation: √4×top surface area÷π (Mathworks 2024). Canopy coverage was measured as the ratio of the vegetative pixel area (from the top-view image) to the total area of the bounding box, which includes both plant and background space (Fig. 5A). Top and side surface area were also measured (Fig 5A, 5B). To the greatest extent possible, plants were carefully removed and promptly returned to the hydroponic system to minimize any adverse effects on plant growth.

### Photosynthesis Analysis

CO_2_ assimilation rates and stomatal conductance of all treatments were measured in fully expanded young leaves using a portable photosynthesis system (LI-6400XT, LI-COR Biosciences, Lincoln, NE, USA) with a clear top chamber, following the methods of Qu et al. (2025) and Yoshiyama et al. (2024), within their respective treatment environments using the same 5000 K W lights and FR lights as during plant growth. 4 weeks into the experiment, the steady-state photosynthetic rate (A) and stomatal conductance (g_s_) were measured in five randomly selected plants grown under W light.

Measurements were first taken under 300 μmol m^-2^ s^-1^ of W light for 5 minutes. FR light was then added, and once A and g_s_ stabilized, they were recorded again under W+FR light for 5 minutes.

The same procedure was applied to five randomly selected plants grown under W+FR light at the 4-week stage: initial measurements were taken under 300 μmol m^-2^ s^-1^ of W light for 5 minutes, then FR light was supplemented. Once A and g_s_ stabilized, they were recorded again under W+FR light for 5 minutes.

### Ascorbic Acid and Pigment Analysis

For lettuce quality analysis at the end of week 5, the contents of ascorbic acid (vitamin C), total chlorophyll *a+b,* total carotenoids, and anthocyanin were measured with 4 to 10 plants measured per treatment (Levine et al. 2023; Hayashi et al. 2024). We selected week 5 for measurements to ensure consistent sampling and reproducibility, as plants at week 6 risked becoming impractically large and less representative of typical consumption stages across repeated trials. For ascorbic acid analysis, two fully expanded, healthy mature leaves were harvested and thoroughly grounded with a pestle and mortar. The juice was pipetted into a 2 mL vial using water for sample dilution. The ascorbic acid in each treatment was quantified by using Reflectoquant^®^ Ascorbic Acid Test strips (Sigma-Aldrich Canada Co., Ontario, Canada) together with an RQ Flex^®^ plus reflectometer instrument reader (Merck Darmstadt, Germany) (Furuta et al. 2025).

For anthocyanin, chlorophyll, and carotenoid analysis, four 6.4 mm diameter (32 mm^2^ holes) disks were punched in on large, fully expanded leaf blades in the darkest region areas that avoided the midrib. We carefully addressed potential anthocyanin variability by collecting all leaf disks from the upper ∼30% portion of the outer edge region of leaves, where anthocyanins most prominently accumulated. Within this region, samples were randomly collected to avoid positional bias and to ensure representative sampling. The mass of the 4 leaf disks was immediately weighed to also assess pigment, anthocyanin and ascorbic acid concentrations on leaf mass basis. For anthocyanin content index (ACI), 1.5 mL of 50:45:5 volume of water, methanol, and acetic acid was added according to Laby et al. (2000) to the 4 leaf disks and placed in a 2 mL vial with a homogenizer bead. This was subsequently shaken at 1100 rpm for 18 minutes in a bead-type crushing device (BMS-A20TP Shake Master, Biomedical Science Inc, Tokyo, JPN). The sample vial was subsequently centrifuged for 2 min, and the supernatant was used for the analysis.

Absorbances at 530.0 and 657.0 nm were measured using a spectrophotometer (UV-1280, Shimadzu Corporation, Kyoto, Japan), and the relative values of anthocyanin content were determined (Laby et al., 2000).

For chlorophyll and carotenoid analysis, 1.5 mL of 80% acetone was added to the 4 leaf disks and placed in a 2 mL vial along with a homogenizer bead. This was subsequently shaken at 1100 rpm for 6 minutes in the same crushing device. The sample vial was subsequently centrifuged for 2 min. Absorbances at 750.0, 663.2, 646.8 and 470.0 nm were measured and the chlorophyll and carotenoid contents were determined using equations from (Lichtenthaler, 1987).

### Gene Expression Analysis

For the anthocyanin biosynthetic gene expression analysis, leaves of 5-week-old plants were sampled when lights were on for 15 hours into their 16-hour photoperiod. The anthocyanin biosynthetic genes including *CHS* (chalcone synthase), *F3H* (flavanone 3-hydroxylase), *DFR* (dihydroflavonol 4-reductase), *ANS* (anthocyanidin synthase), and *UFGT* (UDP-glucose: flavonoid 3-O-glucosyltransferase) were analyzed (Table S1). Total RNA was extracted using an RNeasy Plant Mini Kit (QIAGEN N.V, Hilden, Germany) and reverse transcribed by PrimeScript™ RT reagent Kit with gDNA Eraser (Takara Bio, Japan). RT-qPCR assay was performed with TB Green® Premix Ex Taq™ II (Takara Bio, Japan) using on a Step One Plus Real-Time PCR System (Thermo Fisher Scientific, USA). Eight biological replicates were applied for each gene. *Actin (*ACT, AB359898) was used as an internal control to normalize different samples. The primer sequences used in this analysis are listed in Table S1.

### Statistical Analysis

The experiment followed a randomized complete block design, with 5-10 experimental units for each of the 4 treatments. Plant numbers per harvest varied due to scheduled changes in planting density. In weeks 1–2, we propagated 300 plants, allowing for over 20 viable samples, although we limited sampling to 10 plants. In weeks 3–4, with 100 plants per treatment we maintained a sample size of 10. By weeks 5–6, density dropped to 30 plants per treatment limiting sampling to 5–7 plants while avoiding non-well represented plants on the edges. The experiment was repeated over time for a total of four experimental runs. Analysis of the data were analyzed using JMP Pro 15 software, developed by the JMP business unit of the SAS Institute, using the Tukey–Kramer Honest Significant Difference (HSD) test at *P* = 0.05 to determine significant differences among measured parameters based on light treatment.

## Results

### Plant Growth

The W+FR treatment significantly increased the top canopy surface area (Fig. 5C, 5D), side canopy surface area (Fig. 5E, 5F), and canopy equivalent diameter (Fig. 5K, 5L) at weeks 3, 4, and 5 compared to the W and W to W+FR treatments. These increases in top canopy surface area and canopy equivalent diameter are also visually apparent in the representative images (Fig. 2A), while changes in side canopy surface area can be observed in the representative images (Fig. 2B). At week 3, the W treatment exhibited greater canopy coverage compared to the W+FR treatment; however, by weeks 4 and 5, canopy coverage was similar across all treatments.

**Figure 2:**
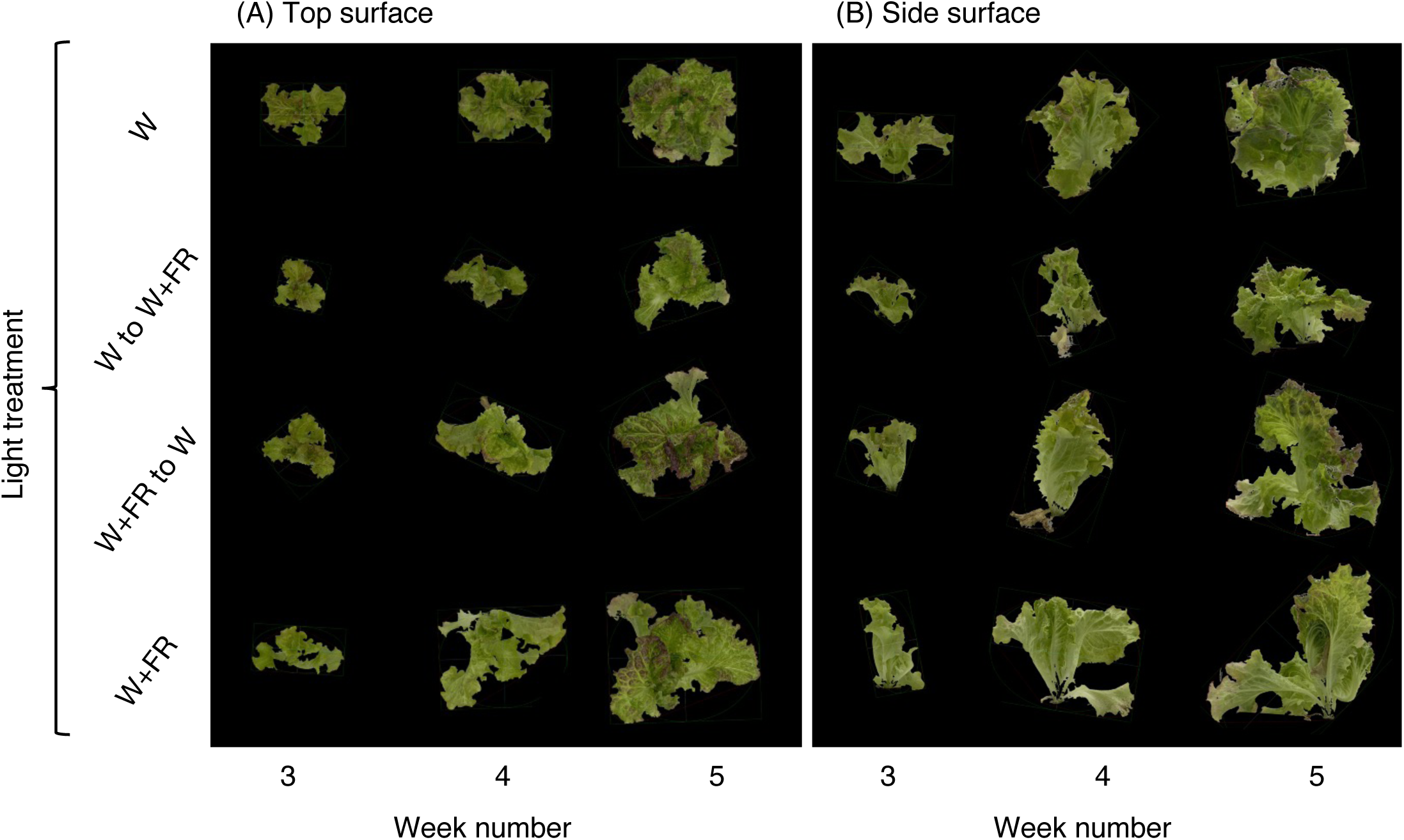
(A) Top surface and (B) side surface views of representative red leaf lettuce grown under continuous white light (W), W for 4 weeks followed by W plus far-red (W+FR), W+FR followed by W, and continuous (W+FR) measured 3, 4, and 5 weeks after respective treatments. The images depict the morphological differences between the four lighting treatments, highlighting the increased leaf expansion and more robust canopy development in the W+FR treatment compared to the W treatment.

Regarding destructive harvest parameters, W+FR resulted in increased shoot fresh and dry weight across most weeks relative to W treatments (Fig. 3A, 3B, 3C and 3D). In week 1, the W+FR treatment resulted in greater dry weights (4.5 ± 0.3 mg) compared to the W treatment (3.5 ± 0.2 mg) but not statistically different fresh weight (Fig. 3A and 3C). In week 2, The W+FR treatment had significantly higher fresh weight (599.5 ± 15.3 mg) but similar dry weight relative to the control (fresh weight: 468.5 ± 16.4 mg) (Fig. 3A and 3C). The differences became more pronounced starting from week 3 with fresh and dry weights (Fig. 3A, 3B, 3C and 3D). As the experiment progressed, plants under the W+FR treatment consistently exhibited the highest dry and fresh weights, while those under the control (W) produced the lowest (Fig. 3A, 3B, 3C and 3D). Interestingly, the W+FR to W treatment had a similarly high fresh weight (113.2 ± 5.2 g) as the W+FR treatment (121.3 ± 5.2 g) by week 6 (Fig. 3B). The treatments where plants were switched from W to W+FR (6.2 ± 0.5 g) or from W+FR to W (5.9 ± 0.3 g) yielded intermediate dry weight by week 6 (Fig. 3D). Although the growth promoting effects of FR light have been reported, few studies have assessed these effects under commercially relevant production conditions. Here, we conducted an experiment using the facility, nutrient management system, and cultivation equipment of a commercial PFAL company in Japan currently operating four active facilities. Plants were initially grown at 1847 plants per m² and transplanted to 314 and 63 plants per m² following standard commercial density reduction workflows. This dynamic adjustment mirrors real-world practices to optimize production. Our findings provide new insights into how FR supplementation affects lettuce biomass and morphology across different density stages as well as anthocyanin biosynthetic gene expression, quality metrics, and photosynthetic parameters. Statistical analysis confirmed that these differences were significant, as indicated by the distinct groupings identified by Tukey-Kramer HSD test (Figs. 2 and 3).

**Figure 3.**
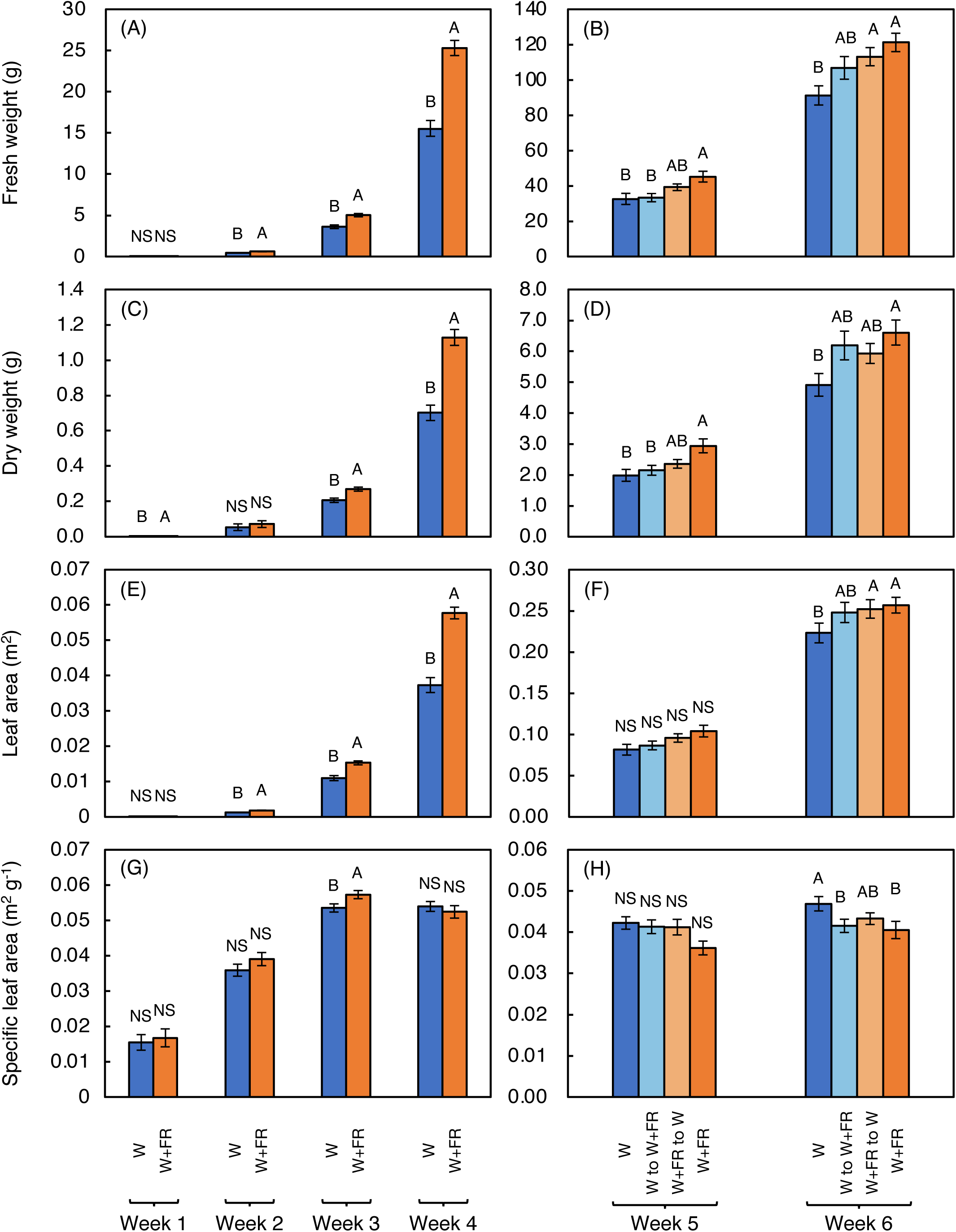
(A) Fresh weight of red leaf lettuce shoots under different lighting treatments from weeks 1 to 4. (B) Fresh weight of red leaf lettuce shoots under different lighting treatments from weeks 5 and 6. (C) Dry weight of red leaf lettuce shoots under different lighting treatments from weeks 1 to 4. (D) Dry weight of red leaf lettuce shoots under different lighting treatments from weeks 5 and 6. (E) Leaf area of red leaf lettuce under different lighting treatments from weeks 1 to 4. (F) Leaf area of red leaf lettuce under different lighting treatments from weeks 5 and 6. (G) Mean specific leaf area of red leaf lettuce shoot from week 1 to 4. (H) Mean specific leaf area from week 5 and 6. Bars represent standard errors (n = 5–10). Different capital letters indicate statistically significant differences between treatments within each time point, as determined by Tukey’s HSD test (p ≤ 0.05).

Regarding leaf area, the W+FR treatment generally promoted greater leaf expansion compared to the control (W), particularly in the early weeks (weeks 2-4), with the most notable difference observed in week 4 (Fig. 3E). As the plants matured from Weeks 5 to 6, the leaf area across all treatments became similar (Fig. 3F). Regarding specific leaf area (SLA), all treatments were similar across all weeks except in week 3 in which W+FR treatment had significantly increased SLA relative to the control. These results indicate that FR supplementation initially enhances leaf area and SLA (in week 3), but these effects become less pronounced as the lettuce plants reach full maturity. Tukey-Kramer HSD test supported these observations, confirming the significant differences (Fig. 3).

### Photosynthetic Parameters

Regarding CO_2_ assimilation rates, leaves acclimated to W and measured under W+FR had the highest CO_2_ assimilation rates, significantly exceeding that of FR-acclimated leaves measured under either W or W+FR (Fig. 4C). In both the W treatment and the treatment where plants were switched from W+FR to W, CO_2_ assimilation rates were enhanced when FR light was supplemented during measurement, compared to when it was not provided. There was no statistically significant difference among the stomatal conductance among all treatments (Fig 4D).

**Figure 4.**
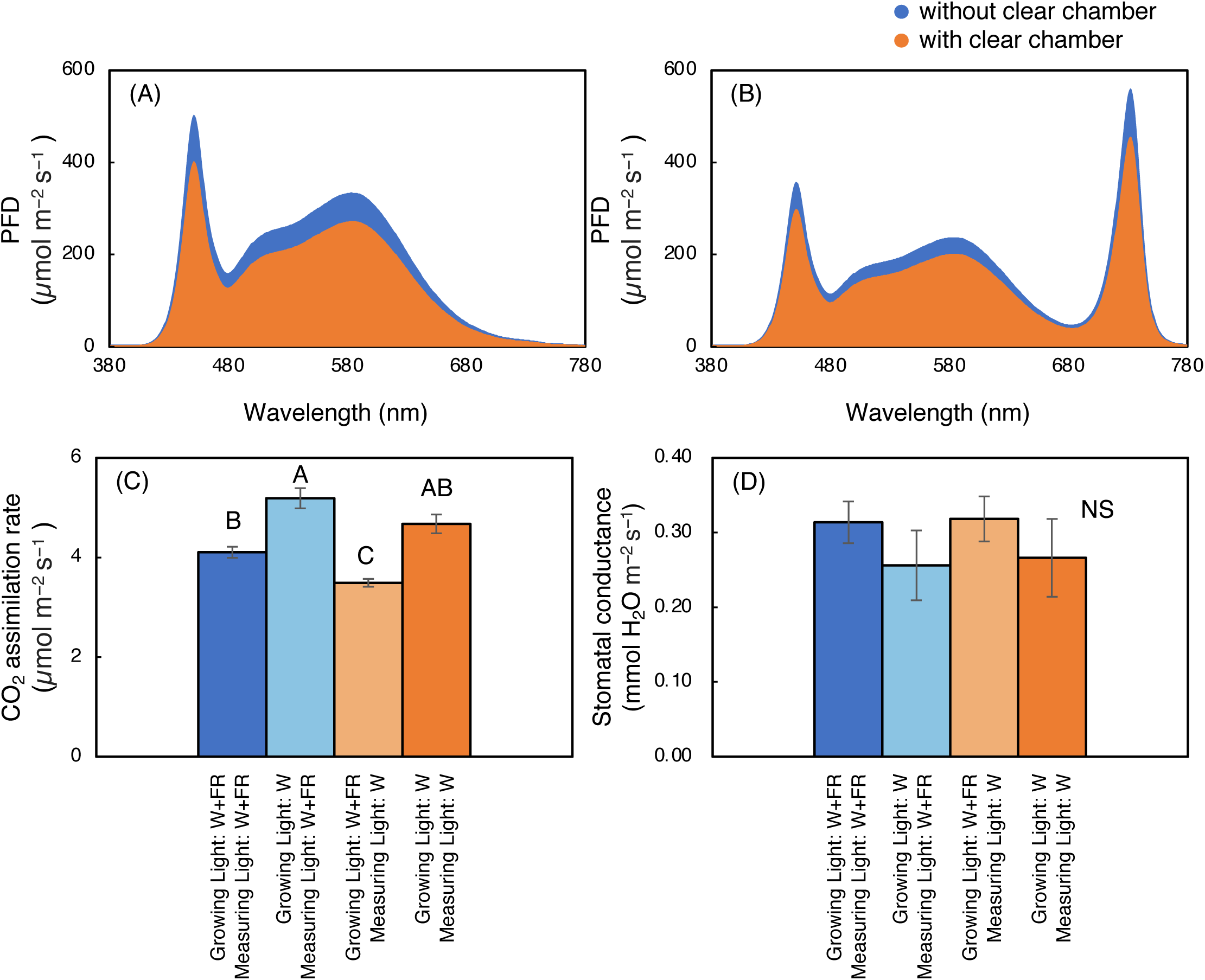
Red leaf lettuce measurements taken 4 weeks after seeding of treatments 1 to 4 of (A) CO_2_ assimilation rate, (B) stomatal conductance, (C) ETR II, (D) ETR I, and (E) ETR I to ETR II ratio. Bars are standard errors (n = 4-5).

Regarding emitted light and leaf absorptance, FR light (700–750 nm) was absorbed substantially less than PAR (400–700 nm) photons in both red and green leaves (Table 2 and Fig S2). While absorption within the 400–700 nm range exceeded 64% in red leaves and 53% in green leaves, total FR absorption was only 7.6% and 6.3%, respectively. Notably, the longer FR waveband (715–750 nm) had the lowest absorption, with red leaves absorbing just 4.1% and green leaves 3.1%. These results indicate a marked drop in absorptance beyond 715 nm. Thus, the photosynthetic contribution of FR photons especially those above 715 nm may be limited due to lower absorptance.

**Table 2.** Incident Light and Leaf Absorptance (µmol m^-2^ s^-1^ and %) Across the 400–750 nm Spectrum in Red and Green Leaves.

| **Color** | **Waveband**<br>**(nm)** | **Incident**<br>**Light**<br>**($\mu\text{mol m}^{-2} \text{s}^{-1}$)** | **Red Leaf**<br>**absorptance**<br>**(%)** | **Green Leaf**<br>**absorptance**<br>**(%)** |
| --- | --- | --- | --- | --- |
| Blue | 400–500 | 88.1 | 90.5 | 86.3 |
| Green | 500–600 | 136.9 | 75.4 | 51.6 |
| Red | 600–700 | 76.4 | 74.4 | 69.6 |
| Far-Red | 700-715 | 11.6 | 29.3 | 25.9 |
| Far-Red | 715-750 | 71.8 | 4.0 | 3.1 |
| **Total** |  |  |  |  |
| Total Far-Red | 700-750 | 83.4 | 7.6 | 6.3 |
| Total Photon<br>Flux Density | 400-750 | 385 | 64.0 | 53.3 |

Table 3 shows that adding FR light (700–750 nm) slightly increases total absorbed quanta beyond the 400–700 nm baseline, but the increase is modest with only about 2.7% for red leaves and 2.6% for green leaves. Under constant photon flux density (PFD) substitution, overall absorption drops to ∼78%, highlighting that FR is absorbed less efficiently than PAR. When absorbed quanta are matched (ideal conditions), FR inclusion is equal to total absorption compared to PAR alone.

**Table 3.**
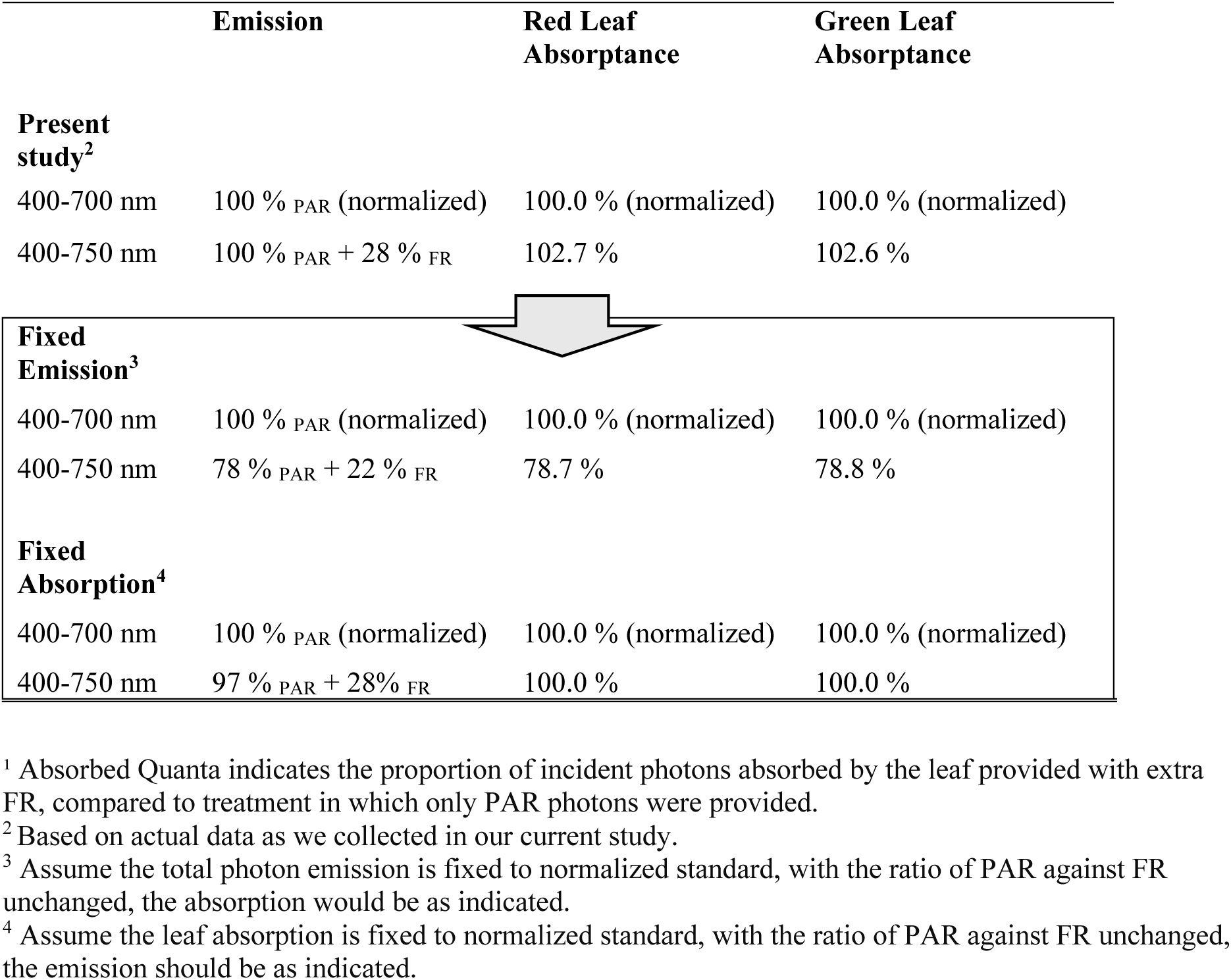
Absorbed Quanta^1^ Under Different Treatments.

|  | **Emission** | **Red Leaf Absorptance** | **Green Leaf Absorptance** |
| --- | --- | --- | --- |
| **Present study<sup>2</sup>** |  |  |  |
| 400-700 nm | 100 % PAR (normalized) | 100.0 % (normalized) | 100.0 % (normalized) |
| 400-750 nm | 100 % PAR + 28 % FR | 102.7 % | 102.6 % |
| **Fixed Emission<sup>3</sup>** |  |  |  |
| 400-700 nm | 100 % PAR (normalized) | 100.0 % (normalized) | 100.0 % (normalized) |
| 400-750 nm | 78 % PAR + 22 % FR | 78.7 % | 78.8 % |
| **Fixed Absorption<sup>4</sup>** |  |  |  |
| 400-700 nm | 100 % PAR (normalized) | 100.0 % (normalized) | 100.0 % (normalized) |
| 400-750 nm | 97 % PAR + 28% FR | 100.0 % | 100.0 % |

<sup>1</sup> Absorbed Quanta indicates the proportion of incident photons absorbed by the leaf provided with extra FR, compared to treatment in which only PAR photons were provided.
<sup>2</sup> Based on actual data as we collected in our current study.
<sup>3</sup> Assume the total photon emission is fixed to normalized standard, with the ratio of PAR against FR unchanged, the absorption would be as indicated.
<sup>4</sup> Assume the leaf absorption is fixed to normalized standard, with the ratio of PAR against FR unchanged, the emission should be as indicated.

### Plant Form - Canopy Architecture

The impact of FR light on plant morphology was assessed by EasyPCC V2 software image analysis from top and side perspectives (Fig. 2 and Fig 5A and 5B). Regarding canopy surface area, the W+FR treatment consistently resulted in the largest top and side canopy surface area (top surface: 1.64 × 10⁶ ± 1.22 × 10⁵ pixels (px), side surface: 3.68 × 10⁶ ± 3.6 × 10⁵ px at 35 days) compared to the W treatment (top surface: 1.06 × 10⁶ ± 1.73 × 10⁵ px, side surface: 2.26 × 10⁶ ± 4.5 × 10⁵ px at 35 days) across each week, indicating that continuous FR exposure promotes significant lateral growth and canopy expansion (Fig. 5C, 5D, 5E and 5F).

**Figure 5.**
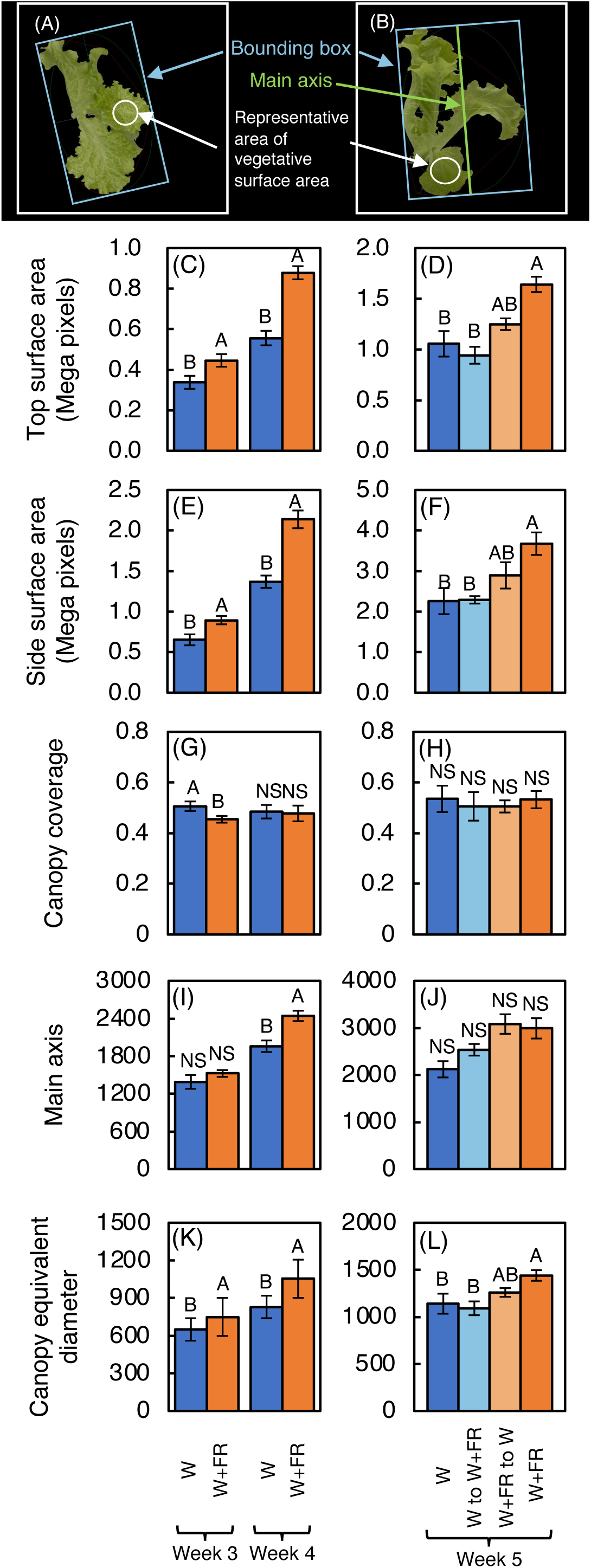
(A) Representative image of the top canopy and (B) representative side image of red leaf lettuce, with explanations of measured parameters. (C) Top surface area from weeks 3 and 4. (D) Top surface area from week 5. (E) Side surface area from weeks 3 and 4. (F) Side surface area from week 5. (G) Canopy coverage (the top canopy surface area, calculated as the actual number of vegetation pixels divided by the area of the bounding box) from weeks 3 and 4. (H) Canopy coverage from week 5. (I) Main axis from the side (a general reference of approximate plant height) from weeks 3 and 4. (J) Main axis from week 5. (K) Canopy equivalent diameter (the diameter of the circle whose area is the same as the pixel surface area of vegetative tissue from the top canopy) from weeks 3 and 4. (L) Canopy equivalent diameter from week 5. Bars represent standard errors (n = 5). Capital letters indicate significant differences by Tukey’s HSD test (5% level of significance).

Significant differences in canopy coverage, defined as the surface area of actual vegetation pixels divided by the bounding box area, were also found. On week 3, the W treatment had the highest canopy coverage, significantly outperforming the continuous W+FR treatment (Fig. 5G). However, as the experiment progressed, the differences in canopy coverage between the treatments diminished, with all treatments showing similar canopy coverage on weeks 4 and 5 (Fig. 5G and 5H). This suggests that while initial differences in canopy coverage may exist, the plants’ overall coverage tends to equalize over time.

The main axis, an indicator of plant height, also varied significantly across the different treatments (Fig. 5I). By week 4, the W+FR treatment had significantly increased height relative to the W treatments (Fig. 5I). However, on weeks 3 and 5, there were no significant differences in the main axis across all light treatments (Fig. 5I and 5J).

Regarding the canopy equivalent diameter, which is the diameter of a circle with an area equivalent to the contour area of the canopy, the W+FR treatment consistently resulted in the largest diameters on all weeks (Fig. 5K and 5L). This trend was evident from the earliest measurements on week 3, where W+FR significantly outperformed the W treatment. Over time, the W+FR treatment maintained its dominance, resulting in the largest canopy diameters by the final measurement on week 5 (Fig 5L). The W treatment, on the other hand, resulted in the smallest diameters, indicating that white light alone does not promote the same level of canopy expansion growth as FR-supplemented treatments (Fig 5K and 5L). The intermediate treatments W to W+FR showed canopy equivalent diameters statistically smaller than W+FR treatment by week 5 and the W+FR to W showed canopy equivalent diameters in between the W and W+FR treatments by week 5 (Fig. 5L).

### Photochemical and Gene Expression Analysis

Regarding chlorophyll *a*+*b* concentration, the control (W) consistently resulted in the greatest chlorophyll content, suggesting that this particular 5000 K W spectral treatment alone is optimal for maximum chlorophyll synthesis and accumulation relative to the other lighting treatments in this study (Fig. 6A and 6B). The W to W+FR treatment had the lowest chlorophyll levels for both per unit area and per unit weight bases, indicating that introducing FR later in the growth cycle might reduce chlorophyll content. The W+FR to W treatment also had significantly lower chlorophyll per unit weight (Fig. 6B).

**Figure 6.**
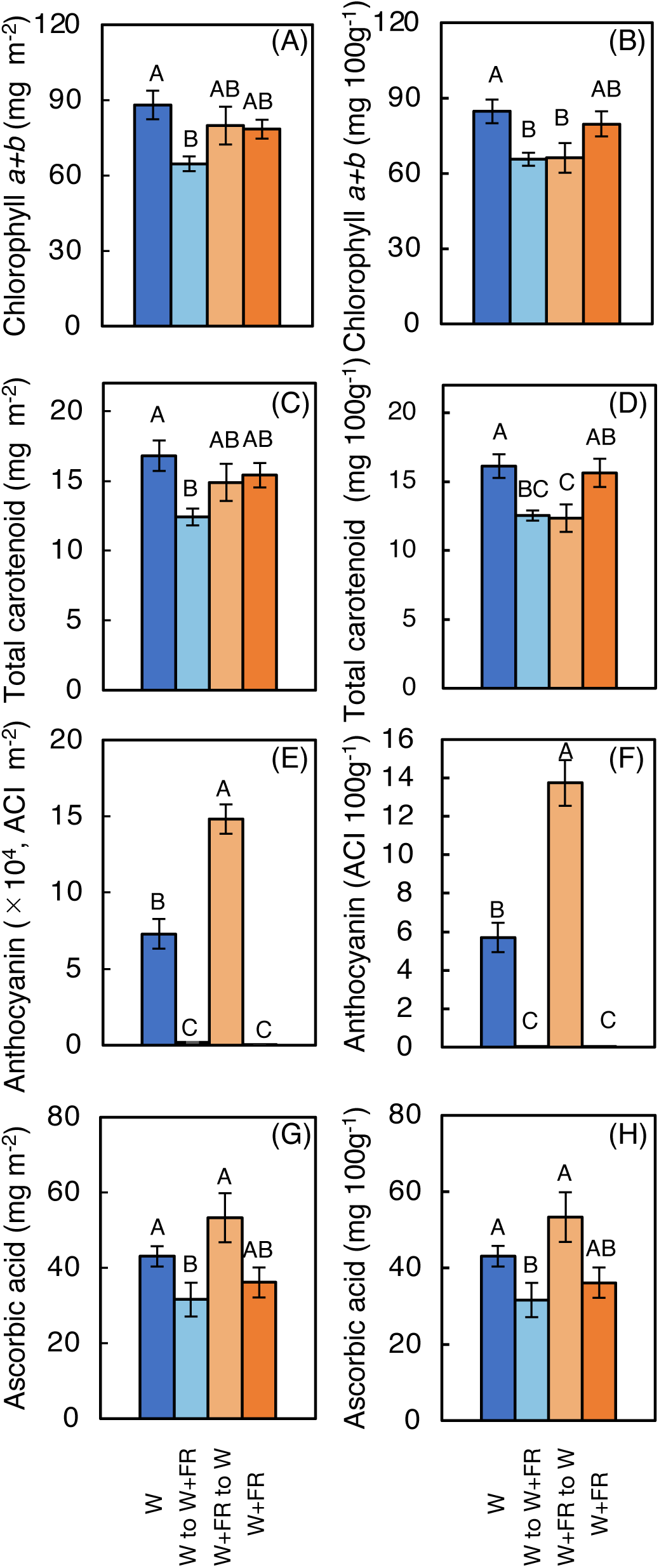
(A) Red leaf lettuce total chlorophyll a+b concentrations 5 weeks after seeding with all treatments with concentration per m^2^ and (B) concentration per 100 grams. (C) Lettuce total carotenoid concentrations 5 weeks after seeding with all treatments per m^2^ and (D) on a total carotenoid concentration per 100 grams. (E) is anthocyanin content index (ACI) in red leaf lettuce 5 weeks after seeding with all treatments on a per m² basis and (F) ACI per 100 grams. (G) is ascorbic acid concentration 5 weeks after seeding with all treatments per m² and (H) ascorbic acid concentration per 100 grams. Bars are standard errors (n = 4-10). Different letters indicate statistically significant differences between treatments, as determined by Tukey’s HSD test (p ≤ 0.05).

Ascorbic acid content was also measured to assess the nutritional quality. The control treatment W and W+FR to W treatments had the highest ascorbic acid contents, indicating that W light alone or when applied at the later-stage is most conducive to maintaining or enhancing this antioxidant in lettuce leaves (Fig. 6G and 6H). In contrast, the W to W+FR treatment resulted in the lowest ascorbic acid content, suggesting that FR supplementation, particularly when applied later in the growth cycle, may diminish the accumulation of ascorbic acid (Fig. 6G and 6H). The W+FR treatment showed intermediate ascorbic acid concentrations, indicating that the timing of FR exposure plays a critical role in influencing this nutrient’s concentration (Fig. 6G and 6H).

The relative expression levels (/ACT) of five key anthocyanin biosynthetic genes— *ANS*, *CHS*, *DFR*, *F3H*, and *UFGT*—were also measured (Fig. 7). Analysis of the data suggests that the W and the W+FR to W treatments exhibited significantly higher expression levels of *ANS*, *DFR*, and *F3H* compared to the continuous W+FR treatment and W to W+FR treatments (Fig. 7). Since the gene expression analysis was measured at the week 5 stage, it is important to note that the W+FR to W treatment had already transitioned to the W treatment for one week and was no longer receiving FR supplementation at the time of measurement. In contrast, *CHS* and *UFGT* gene expressions showed no significant differences across any of the treatments, indicating that these genes were less responsive to the lighting conditions tested (Fig. 7).

**Figure 7.**
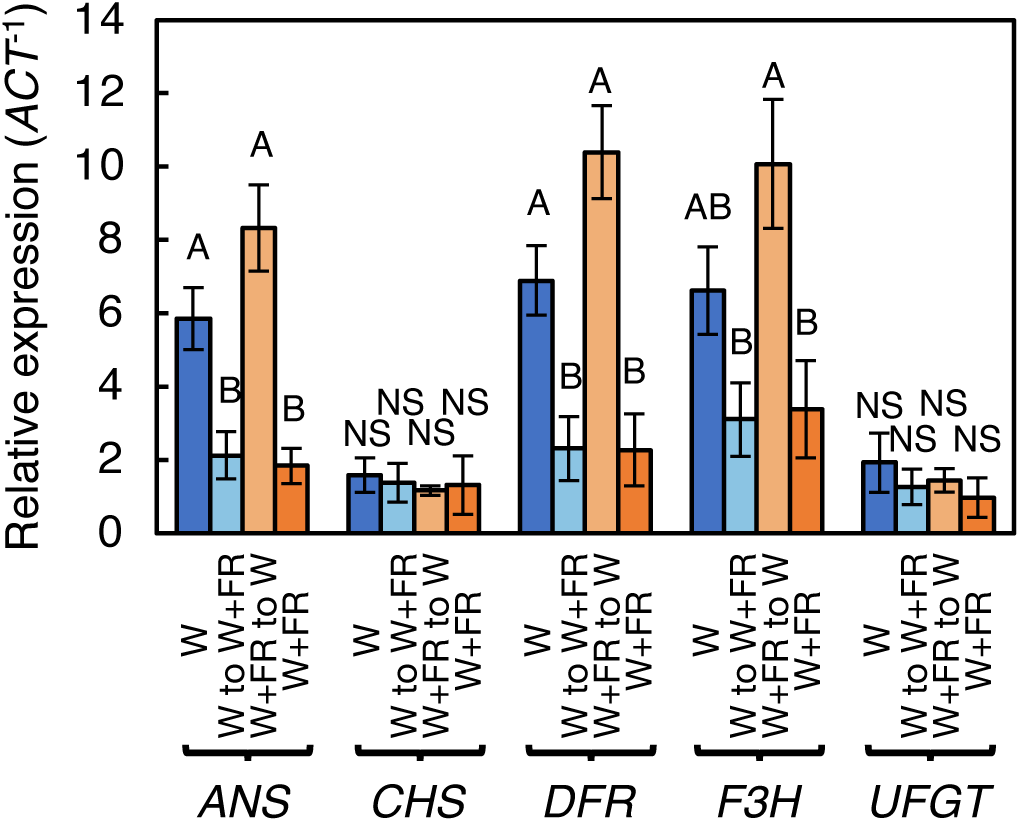
Anthocyanin biosynthetic gene expression analysis of fresh leaf tissue from red leaf lettuce 5 weeks after seeding under different lighting treatments: (1) white light (W), (2) white light followed by far-red (W to W+FR), (3) far-red followed by white light (W+FR to W), and (4) continuous far-red (W+FR). Capital letters indicate statistically significant differences between treatments, as determined by Tukey’s HSD test (p ≤ 0.05). Bars represent standard errors (n = 10).

### General Evaluation

Lastly, to summarize all the results, the radar plot illustrates the comparative effects of all treatments on key plant growth and phytochemical parameters, including fresh weight, leaf area, chlorophyll *a*+*b*, total carotenoid, anthocyanin, and ascorbic acid concentrations (Fig. 8). The W+FR to W treatment generally demonstrate superior outcomes across all parameters while W+FR treatment leans more towards superior plant growth parameters and compromises on anthocyanin phytochemicals relative to other treatments. The W to W+FR treatments shows a stronger tendency for plant growth parameters over phytochemical synthesis. Lastly, the W treatment leans more towards phytochemical synthesis while sacrificing plant growth (mainly fresh weight) parameters. This analysis underscores the differential impact of dynamic light conditions on both growth and phytochemical profiles.

**Figure 8.**
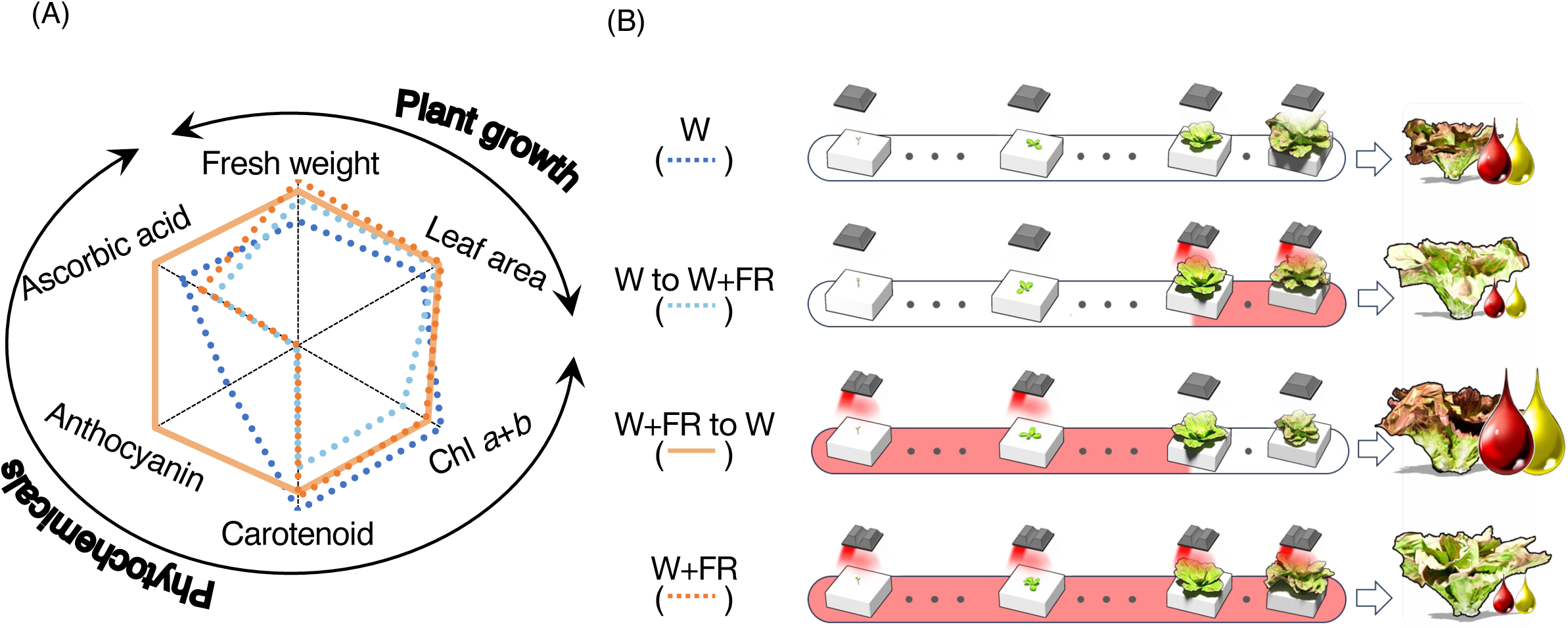
(A) Radar plot among four treatments and (B) the illustration of summary. The radar plot compares the effects of four different treatments: (1) white light (W), (2) white light followed by far-red (W to W+FR), (3) far-red followed by white light (W+FR to W), and (4) continuous far-red (W+FR) on various growth and phytochemical parameters in plants. The parameters measured are: fresh weight, leaf area, chlorophyll a+b content, total carotenoid content, anthocyanin content, and ascorbic acid content. These parameters are categorized into two broad groups: Plant growth (fresh weight and leaf area) and phytochemicals (chlorophyll, carotenoids, anthocyanin, and ascorbic acid). The illustration of summary visually shows the general timeline for each treatment and the light regime transitions, showing how the type and timing of light exposure influenced lettuce growth and phytochemical accumulation.

## Discussion

Light is one of the most important environmental factors determining plant growth and phytochemical concentrations in plants in artificial light plant factories (Kopsell and Kopsell, 2008, Perez-Balibrea et al., 2008; Matsuura, Costa, Yendo, & Fett-Neto, 2013). As red and blue light are efficiently absorbed by chlorophylls, the effects of red and blue radiation on growth and nutritional properties of lettuce have been the focus of numerous past and present studies (Kaiser et al., 2019; Li et al. 2025).

In addition, UV has been reported to enhance phytochemicals, including anthocyanin, carotenoids and chlorophylls (Vergeer et al., 1995, Keller et al., 1998; Ohashi-Kaneko et al., 2007; Tsormpatsidis et al., 2008; Chen et al., 2019; He et al. 2021). Applying blue light radiation at the end of the production cycle can significantly increase anthocyanin concentrations in red-leaf lettuce, improving its coloration and nutritional quality (Kelly et al, 2023, Zhu et al., 2024, Owen et al., 2015). It has also been shown that supplemental FR light can decrease anthocyanin concentration but promote leaf extension and plant growth in lettuce (Gommers et al., 2013; He et al., 2021).

However, there is no universally optimal FR or broader spectral lighting recipe (e.g., 300–800 nm) for plant cultivation in plant factories, as light requirements vary depending on plant species, growth stage, and cultivation objectives (e.g., flowering, vegetative growth, fruit production, postharvest quality) (Barbieri et al., 2023; Thilini Deepashika Perera et al., 2022; Piovene et al., 2015). Furthermore, it has been stated that “every VF [vertical farming] system is different” (Carpineti et al., 2024), and variability exists between research and production modules thus making consolidated findings from different studies not straightforward.

The present work showed that the FR treatment led to an increase in leaf area in lettuce (Fig. 3E and 3F), improving light interception and significantly increasing biomass production (Fig. 3A, 3B, 3C and 3D). However, this was accompanied by a reduction in the localized concentration of anthocyanins (Fig. 6E and 6F). Furthermore, compared to the W treatment, the W to W+FR treatment resulted in a significant decrease in total carotenoid concentrations both on area and weight bases. (Fig. 6C and 6D). However, while total carotenoids per leaf area were significantly lower following the W to W+FR treatment, carotenoids per leaf mass were not significantly different from the W treatment (Fig. 6C and 6D). Thus, there is a potential trade-off between yield and quality.

Although spectrophotometric measurements in Fig. 6E and 6F show significantly higher anthocyanin accumulation than what appears in the representative images in Fig. 2, it is important to note that anthocyanin is highly localized. The sampled leaf disks were taken from the darkest regions with the highest pigmentation and do not reflect anthocyanin levels across the entire plant shown in Figure 2. In contrast, the absence of FR light on week 5, 1 week prior to final harvest increased the accumulation of desirable chemical compounds, including anthocyanins and ascorbic acid, in plants grown under W light 1 week before harvest (Fig. 6 E, 6F, 6G and 6H). Thus, applying FR light only during the early growth phase could improve the plant production, balancing nutritional quality and yield (Fig. 3B and Fig. 6E, 6F, 6G and 6H).

Analysis from the radar (spider) plot data suggests that FR radiation applied during the initial two-thirds of the production cycle (weeks 0–4 of a 6-week cycle) can be strategically utilized to enhance plant growth and phytochemical content in plants grown under W light during the later stages in a commercial PFAL with this cultivar under these growing conditions (Fig. 8).

### FR light enhances plant growth by improving architecture and photosynthesis

The increased biomass accumulation observed with supplemental FR light (Fig. 3A, 3B, 3C and 3D), the increased CO_2_ assimilation rates with W acclimated leaves measured under W+FR (Fig. 4C), and the expanded leaf canopy area (Fig. 5) suggest that FR supplementation enhances plant growth through a combination of increased canopy expansion and CO_2_ assimilation. Several studies have shown that fluctuating light observed in the natural light environments causes PSI photoinhibition highlighting the potential vulnerability of PSI under changing light conditions (Kono and Terashima 2014; Suorsa et al., 2012; Yamori et al., 2016; Yamori & Shikanai, 2016). The addition of FR to the fluctuating light suppressed PSI photoinhibition via keeping oxidation of P700, the reaction center of PSI (Kono et al., 2017). More recently, it has been reported that FR exerted beneficial effects on photosynthesis in fluctuating light by exciting PSI and accelerating the relaxation of non-photochemical quenching (NPQ) in the PSII antenna system, resulting in increases in photosynthesis (Kono et al. 2020).

On the contrary, Zhen and Bugbee (2020) reported that substitution of FR photons maintained an equivalent canopy quantum yield, thereby suggesting that FR photons can contribute as effectively as PAR photons to photosynthesis when acting synergistically.

However, more recently, Jeong et al. (2024) suggested that under FR light substitution, plant growth may become more dependent on total photon capture rather than photosynthetic efficiency. Jeong et al., (2024) reported a significant reduction in the net photosynthetic rate per unit leaf area when a 20% substitution of FR photons was applied to FR acclimated leaves. In a recent review, Shomali et al. (2025) stated “If such a link exists between the chloroplast and the nucleus, then it raises the opinion that FR perception takes part in driving photosynthesis through the perception of chloroplastic retrograde signals.” This perspective emphasizes that if FR is shown to regulate photosynthesis beyond simple photochemical responses, then its integration into light management strategies could be more foundational than currently understood. However, such a regulatory mechanism between FR perception and photosynthetic enhancement remains to be fully elucidated. Until such a mechanism is conclusively demonstrated, the proposal to regard photons in the 400–750 nm range as equivalent photosynthetic photons, without accounting for leaf absorptance cannot be considered best practice.

It is also noteworthy that independent nonprofit organizations such as Design Lights Consortium (DLC), in its latest Technical Requirements for LED-Based Horticultural Lighting, Version 3.0, maintains its evaluation of photosynthetic photon flux (PPF) and photosynthetic photon efficacy (PPE) within the American National Standards Institute (ANSI) / Illuminating Engineering Society (IES) LM-79 standard of measuring photons between the 400–700 nm range. This standard reflects the well-established definition of PAR as the wavelengths most directly contributing to photosynthesis.

Overall, FR photons fall outside the established PAR spectrum (400–700 nm) and contribute less to the energy that directly drives photosynthetic reactions, resulting in a small effect in a stable, steady-state light environment (Fig. 4). Today, commercial CEA typically relies on stable, high-PAR lighting environments, focusing primarily on achieving PPFD and DLI targets. Nevertheless, with the advent of LEDs and multichannel lighting systems, horticultural lighting technologies are advancing toward more dynamic light management, and insights into PSI stability under varying light conditions may become increasingly relevant to CEA.

FR light influences not only photosynthesis (Fig. 4C) but also plant morphology (Figs. 2, 5C, 5D, 5K, 5L, and 8), exerting significant effects on both physiological and structural aspects of plant growth. The absorption of FR photons influences phytochrome systems rather than chlorophyll, meaning the plant’s response is more focused on structural changes for optimizing light use efficiency rather than boosting photosynthetic output. The present work showed that the supplemental FR treatment led to enlarged leaves (Fig. 2) and expanded canopy size (Fig. 5C and 5D), which enhanced light interception and, together with increased CO_2_ assimilation (Fig. 4C), likely contributed to the significant increase in plant biomass (Fig. 3A, 3B, 3C, and 3D). These results indicate that FR promotes plant growth through both improved photosynthetic performance and morphological changes. These findings are consistent with previous research that showed FR supplementation and substitution, promotes stem elongation and leaf expansion (Eylands and Mattson, 2023; Zhen et al., 2020; Kusuma et al., 2023), which is a response likely to enhance light-foraging capacity (Franklin, 2008), and cell wall-modifying mechanisms are vital regulatory points for control of this elongation responses (Sasidharan et al., 2008).

In conclusion, supplemental FR light effectively promoted plant growth by stimulating both morphological changes and photosynthetic activity. W+FR-acclimated leaves showed increased CO_2_ assimilation rates when measured under W+FR light, although the rates were not as high as those of W-acclimated leaves measured under the same light condition. These results highlight that FR supplementation contributes to enhanced biomass production through its combined effects on plant architecture, light interception, and, to some extent, CO_2_ assimilation under the tested air temperatures and light intensities.

Such architectural changes may be particularly advantageous under high density PFAL planting conditions, where maximizing light interception per unit ground area is essential for optimizing productivity. FR LEDs are also among the most efficient in terms of photon output per unit energy input (µmol J^-1^) due to the lower energy of FR photons, making them valuable for reducing energy consumption in PFAL systems (Kusuma et al., 2020).

### Early-stage FR radiation promotes growth without compromising the final concentrations of functional compounds

While continuous FR supplementation throughout the entire 6 week growth period enhances plant biomass (Fig. 3B and 3D), it results in a decrease in the concentration of phytochemicals, such as anthocyanin and ascorbic acid (Fig. 6E, 6F, 6G and 6H). In contrast, this study revealed that supplementing FR radiation (W+FR) only during the first 4 weeks of growth and followed W results in final plant growth comparable to that observed with continuous FR radiation. W+FR to W also significantly increased anthocyanin concentrations (Fig. 6E and 6F) and maintained greater ascorbic acid concentration relative to W to W+FR and similar ascorbic acid concentrations to W and W+FR treatments (Fig. 6G and 6H). This suggests that the initial exposure to FR followed by a shift to W light may trigger a stress response or a metabolic shift that leads to increased anthocyanin accumulation. Moreover, since the W+FR to W plants were taller and closer to the W lights at the time of measurement, the increased light intensity they received could have further amplified anthocyanin synthesis, explaining the significantly greater concentration compared to the continuous W treatment.

The gene expression analysis revealed that W+FR to W treatment resulted in significantly higher expression levels of key anthocyanin biosynthetic genes (*ANS*, *DFR*, and *F3H*) compared to the continuous W+FR treatment and the W to W+FR treatment (Fig. 7).

Some of these anthocyanin biosynthesis genes expressed in our study are also consistent with ones expressed in Goto et al. (2016). Since the W+FR to W plants were generally taller and closer to the lights, thereby receiving greater light intensity than the W treatment, which may explain why their gene expression was numerically greater than in the W treatment. It is well known that anthocyanins contribute to photoprotection by attenuating excess light, while ascorbic acid mitigates oxidative stress by scavenging reactive oxygen species under high light conditions (Szymańska et al., 2017). While continuous W+FR treatment alone might reduce *ANS*, *DFR*, and *F3H* gene expression, it may still play a crucial role in enhancing gene expression when followed by only W light. Alternatively, FR radiation induced a shade avoidance response in the plants, leading to thinner leaves and a shade-type phenotype as a low light response. Consequently, exposure to W light later could trigger a stress response and thus the plants may have exhibited an increased sensitivity to moderate light conditions (300 μmol m^-2^ s^-1^), which are not considered low for PFAL lettuce production.

Our results specifically demonstrate that after four weeks of W+FR treatment, shifting to W light alone for one additional week is sufficient to significantly increase anthocyanin accumulation. This W+FR to W treatment maintains fresh mass comparable to continuous W+FR treatment while significantly enhancing anthocyanin pigmentation. It provides a practical light management strategy that growers can implement during the final production phase to stimulate leaf pigmentation through the plant’s natural stress response to spectral changes, without the need to invest in multichannel LED horticulture systems. Simply transplanting crops to a different section of a PFAL for increased spacing with a preset fixed spectrum of 5000K white is sufficient to achieve this effect for ‘Red Fire’ red leaf lettuce.

These findings have important implications for the commercial production of red leaf lettuce, where leaf color is often associated with perceived quality (Owen et al., 2015); however, while the W+FR to W treatment significantly increases anthocyanin levels relative to other light regimes, our study does not assess whether the absolute pigmentation achieved meets consumer expectations or commercial marketability thresholds. We therefore focus our conclusions on the physiological and production aspects of anthocyanin enhancement.

In addition to the W+FR to W treatment demonstrated in our experiment, other techniques have been reported to stimulate anthocyanin production. These include but are not limited to end-of-production root-zone temperature control (Levine et al., 2023, Hayashi et al., 2024), nutrient stress (Jezek et al., 2023), supplemental blue (B) light application (Kelly and Runkle, 2023, Zhu et al., 2024), controlled exposure to 310 nm UV-B (Goto et al., 2016), and 315-399 nm exposure to UV-A (Kelly and Runkle, 2023). Each of these approaches applies a different abiotic stressor to trigger secondary metabolite production, including anthocyanins, in leafy greens.

However, not all commercial PFALs, such as the commercial PFAL used in our study, are equipped with multichannel lighting systems with independently dimmable channels such as UV-A, UV-B, B, and W, root-zone temperature control systems, or targeted individual macronutrient management capabilities. Consequently, the range of techniques commercially available for controlling anthocyanin accumulation may be constrained by the technological infrastructure of each PFAL.

Although many previous works report that increased FR supplementation decreased phytochemical concentrations, including anthocyanin, carotenoid, and ascorbic acids in lettuce (Li and Kubota 2009, Stutte and Edney 2009, Zou et al., 2023), our study showed that early-stage FR radiation promotes growth without affecting the final levels of functional compounds. Understanding the optimal balance between FR exposure and pigment accumulation is essential for producing visually appealing lettuce with high anthocyanin content, particularly in CEA systems where light quality can be precisely managed.

### Ecological significance

The ecological significance of this study lies in recognizing that plants are evolutionarily adapted to dynamic spectral environments shaped by seasonal shifts, climate, and altitude which are conditions that are currently difficult to replicate in plant factories.

Studies by Siriwardana and Kume (2025) demonstrate how spectral quality varies with airmass and across months with R:FR ratios being the greatest in the summer and lower in the winter. These spectral changes can influence plant growth. This contextualizes the finding from Jeong et al. (2024) in which high FR and high temperature reduced photosynthetic rates and biomass in lettuce which is a cool season crop poorly suited to such environments. It may also explain why Skabelund et al. (2025) observed minimal leaf area expansion in spinach (*Spinacia oleracea*) with increasing FR fractions, even though Kusuma and Bugbee (2023) reported leaf area and other morphological changes from similar FR fraction treatments in strong and contrasting morphological responses in cucumber (*Cucumis sativa* ‘Straight Eight’), green butterhead lettuce (*Lactuca sativa* ‘Rex’).

These discrepancies highlight that plant responses to spectral cues are species-specific and possibly tied to their native environments, whereas the spectral conditions in plant factories are often static and ecologically disconnected from the plants’ natural adaptive context.

### Practical implications

One of the significant findings of this study is the demonstrated effect of FR supplementation under commercially realistic high-density growing environments, where spectral quality, planting density, and resulting canopy light interception collectively influence shoot biomass and pigment accumulation. By performing week-by-week destructive harvests across three practical spacing densities representative of commercial Japanese PFAL setups, this study provides a uniquely detailed dataset capturing changes in shoot biomass, leaf area, and SLA. These results not only validate the practical utility of the W+FR to W light shift for enhancing anthocyanin accumulation without compromising fresh mass but also offer concrete reference data for modeling potential head lettuce yields in NFT based PFAL systems.

Furthermore, the promotion of lateral growth and canopy expansion under FR light, as evidenced by the increased canopy surface area and equivalent diameter (Fig. 5A, 5B, 5C, 5D, 5K and 5L), suggests that plants grown under FR exposure may require more space to avoid overcrowding and ensure optimal light penetration to lower leaves. The findings also indicate that FR exposure can influence canopy architecture in ways that affect planting density. For instance, plants that were exposed to FR light (W+FR) resulted in significantly greater canopy surface area (Fig. 5C and 5D), top, side surface area (Fig. 5E and 5F), and canopy equivalent diameter (Fig. 5K and 5L), relative to the control treatment. This suggests that they might need different planting densities for more efficient growth. Overall, FR exposure may require careful monitoring and potentially increased spacing to prevent excessive shading and competition among plants. The implications for commercial CEA operations are clear: while FR supplementation offers substantial benefits in terms of biomass accumulation (Fig. 3A, 3B, 3C and 3D), these must be balanced with considerations of planting density and spacing. Overcrowding could diminish the benefits of FR supplementation by increasing competition and reducing light penetration, particularly in dense planting arrangements (Jin et al., 2021). Therefore, optimizing light regimes must involve a holistic approach that considers not only the timing and intensity of FR exposure but also the spatial arrangement of plants to maximize overall system efficiency. The vertical space should also be considered since FR tends to elongate the lettuce and cause plants to reach closer to the lights. This suggests that vertical space should be considered when FR is supplemented or substituted in PFAL production. In small multilayer PFALs, the inclusion of FR may also alter air distribution by affecting plant architecture, which influences canopy airflow and an effect highlighted in CFD simulations sensitive to crop drag coefficients (Kang et al., 2024).

It has also been demonstrated that as plant growth progresses, light does not reach the lower part of the canopy, leading to senescence (Boonman et al., 2006; Thomas and Stoddart, 1980). Recent work showed that upward lighting from the base combined with top lighting to reach lower leaves could be applicable to retarding senescence of lower leaves as well as improvement of plant growth (Zhang et al. 2015; Joshi et al. 2017; Saengtharatip et al. 2021; Yamori et al. 2021). In high-density PFAL systems, where maximizing space is critical, the enhanced canopy expansion could lead to increased competition for light (Miao et al., 2023; Kozai 2013), potentially necessitating adjustments in plant spacing to maintain high yield and quality. Thus, in plant factories, optimizing not only plant density but also the wavelength and direction of the irradiated light is recommended for successful cultivation.

## Conclusion and Future Research

Although leaves sometimes contain anthocyanins and other pigments, FR light is absorbed exclusively by chlorophyll. In terrestrial plants, this is chlorophyll *a*, and the absorption spectra of leaves and leaf canopies are well documented. Therefore, in studies analyzing the effects of FR light, it is crucial to consider both the emission spectra of the light source and the absorption spectra of leaves or leaf canopies, and to determine the experimental settings.

Future research may build on the findings of Zhen and Bugbee (2020) by considering a more targeted approach to the application of 700–750 nm FR light. Rather than substituting 1 FR photon for 1 PAR photon, it may be beneficial to tailor FR supplementation based on its relative absorptance and in consideration of species-specific responses and environmental conditions in PFAL systems. Given the lower absorptance of 700–715 nm and especially 715-750 nm FR light relative to PAR wavebands in our ‘Red Fire’ red leaf lettuce, thoughtful experimental design remains critical in photosynthesis studies.

## Supplementary Data

Supplementary data are available at Annals of Botany online and consist of the following: Table S1. List of specific primer sets of the anthocyanin biosynthetic genes used for the gene expression analysis. Figure S1: 3 spacing densities of experiment. Figure S2. Leaf light absorptance of two parts of leaves of ‘Red Fire’ red leaf lettuce.

## Acknowledgments

We acknowledge Hiromasa Kodama for his conscientious technical assistance in helping assist Keiichiro Tanigawa measure photosynthesis under +/- FR in this study. We also acknowledge Akira Ito for his assistance in measuring leaf absorptance.

## Data Availability Statement

The data that support our paper can be requested by contacting the corresponding author.

## Author Contributions

W.Y. and C.P.L. conceived and designed the experiments. C.P.L. supported the set-up of the hydroponic cultivation system. C.P.L. cultivated lettuce hydroponically. C.P.L. mainly performed the experiments and analyzed the data with support from I.T. and W.Y. Y.W. conducted gene expression measurements and analysis. H.K. assisted K.T. with photosynthesis measurements. K.T. analyzed the photosynthesis measurements. Y.Q. helped assist with pigment measurements. W.G advised with the image capturing and analysis. C.P.L. and W.Y. prepared the figures and manuscript with support from K.T., Y.K., I.T., and Y.Q. All authors have read and approved the final version of this manuscript.

