## Supplementary figures and images for "Far-Red Light in Early Growth Stages Boosts Lettuce Biomass and Preserves Anthocyanins"

### fig s1

**Figure S1.**

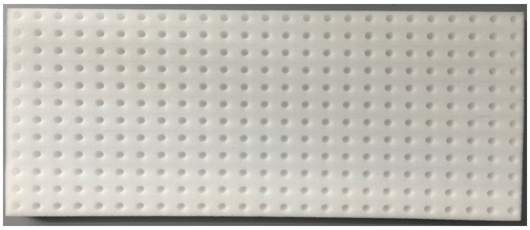

First 2 weeks

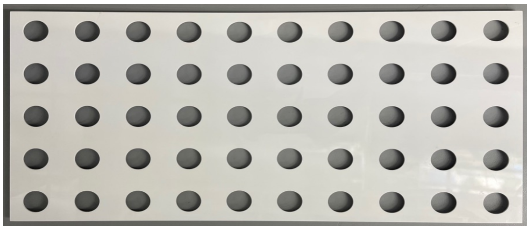

Next 2 weeks

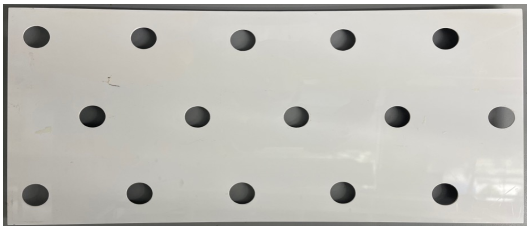

Last 2 weeks
