## Supplementary material for "Far-Red Light in Early Growth Stages Boosts Lettuce Biomass and Preserves Anthocyanins": fig s2

**Figure S2.**

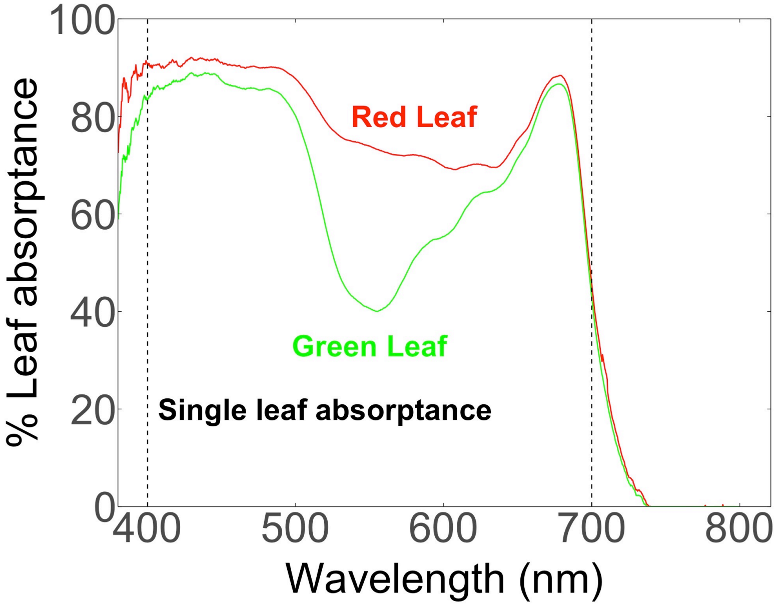


**Figure S2.** Leaf light absorptance of two parts of leaves of red leaf lettuce. The space between the two dotted vertical lines indicates the PAR region (400-700 nm).
