## Supplementary material for "Far-Red Light in Early Growth Stages Boosts Lettuce Biomass and Preserves Anthocyanins": table s1

Table S1. The specific primer sets of the anthocyanin biosynthetic genes used for the gene expression analysis.

| Gene name | Accession number | Product length | Name | Primer sequence (5′−3′) |
|---|---|---|---|---|
| *ACT* | AB359898 | 113 bp | LsACT-F01 | TGGTAGGTATGGGCCAGAAA |
| | | | LsACT-R01 | GTCATCCCAGTTGCTCACAA |
| *CHS* | AB525909 | 169 bp | LsCHS-F02 | GGAGGTGGGGCTAACTTTTC |
| | | | LsCHS-R02 | GAGCTCCACCTGGTCCAATA |
| *F3H* | AB525910 | 210 bp | LsF3H-F02 | CTACTCAAGGTGGCCCGATA |
| | | | LsF3H-R02 | AATGTGAGATCGGGTTGAGG |
| *DFR* | CV700105 | 105 bp | LsDFR-F01 | GGGAATGAGGGAGTGATGAA |
| | | | LsDFR-R01 | ATTGGCAGAAAAAGCAFGCAT |
| *ANS* | AB525912 | 117 bp | LsANS-F01 | CTCCCCACCATCGACTTAAA |
| | | | LsANS-R01 | ATGGTTGACGAGATGCATGA |
| *UFGT* | AB525911 | 203 bp | LsUFGT-F02 | AAGAGACCAGAACCCCGTTT |
| | | | LsUFGT-R02 | AGCTCCAATGCTCTCCGATA |
